# Gene therapy prevents disease and death from non-ketotic hyperglycinemia

**DOI:** 10.1101/2025.03.26.645560

**Authors:** Alejandro Lopez-Ramirez, Adviti Bali, Md Suhail Alam, Prasad Padmanabhan, Shaun Calhoun, Caroline Bickerton, Ana L. Flores-Mireles, Kasturi Haldar

## Abstract

Genetic defects in glycine decarboxylase (GLDC) cause non-ketotic hyperglycinemia (NKH), a rare and frequently fatal neurometabolic disease, which lacks FDA-approved therapies. We characterized CRISPR Cas9-edited humanized mice expressing a prevalent clinical mutation after administration with a single intraperitoneal dose of a novel recombinant of adeno-associated viral vector 9 expressing GLDC (rAAV9-GLDC). Long term biological activity of rAAV9-GLDC was first validated by assessment of its systemic efficacy over five and ten months in mice. Access of rAAV9 to the brain was confirmed by tracking green fluorescent protein (GFP) after a single intraperitoneal dose of rAAV9-GFP. Over five months, control ‘mock’ treated GFP-mice showed reduction in astrocytes but not microglia, oligodendrocytes or neurons in the brain. 37% of these animals suffered long term neurological disease and/or death. rAAV9-GLDC boosts astrogenesis without triggering an inflammatory response and confers 100% protection against disease progression and fatality due to NKH.

## Introduction

The Glycine Cleavage System (GCS) is a major pathway of glycine catabolism and a key mitochondrial metabolic process in vertebrates ^1^. Genetic defects in the GCS cause a rare neurometabolic disorder, non-ketotic hyperglycinemia (NKH), diagnosed by blood-glycine elevation and genetics ^2, 3^. Clinical disease presentation can be highly heterogeneous and classified as either severe or attenuated ^4–6^. In newborns, severe NKH presents with seizures, hypotonia, lethargy, and respiratory apnea, which may require ventilatory support. A substantial portion of affected infants die and those who survive present significant neurological impairment^7^. Patients with mild forms of the disease have also been diagnosed in adulthood ^8, 9^. Present palliative therapies of sodium benzoate and dextromethorphan used to reduce blood glycine levels and seizures can frequently be ineffective: in addition, these compounds must be taken in high and repetitive doses on a daily basis ^10, 11^, suggesting a need for early, curative therapies.

The GCS is prominently expressed in the brain, liver and kidney. Glycine decarboxylase (GLDC) is the rate limiting step of glycine cleavage. Mutations in *Gldc* account for up to 85% of NKH cases. Recent studies have linked increased GLDC expression to psychosis ^12^, reinforcing its role in affecting neurological disease in the brain. GLDC is distributed across all major regions of the brain^13^. It is highly expressed in astrocytes, primary glial cells that may constitute up to 40% of all brain cells ^14, 15^. In contrast, there are no published reports that microglia (immune cells which make up 5-15% of cells in the brain) express the GLDC protein. Low levels of GLDC have recently been reported in subtypes of neurons ^16^. Data from mice confirms that astrocytes are the major reservoirs of GLDC, microglia are depleted, while neurons and oligodendrocytes express extremely low levels (brainrnaseq.org). Recent studies suggest that *Gldc*-deficient neural precursor cells derived from human induced pluripotent stem cells (iPSCs) give rise to a heterogeneous astrocyte lineage ^17^. However, whether mutations in GLDC affect astrocytes in the brain *in vivo* as well as the consequences of astrocyte dysfunction and corrections thereof, for post-natal, cerebral NKH disease and death remain entirely unknown.

The lack of suitable pre-clinical models has severely hindered both the understanding of the pathophysiology of neurological disease that arises due to deficiency in GLDC as well as the development of new treatments for NKH. The most widely used NKH mouse model has been the gene trap mouse model^18^, that yields post-natal mice with (null) deletion in *Gldc*. This model is associated with high levels of prenatal death, presents glycine elevation and folate depletion but not post-natal pathologies of cerebral and neurological disease, or survival defects, likely because it fails to capture mutation-induced disruption of protein-protein interactions expected to contribute to the vast majority of clinical disease. We undertook large-scale genotype-phenotype studies to evaluate disease severity of over two hundred patient *Gldc* mutations ^6^. This led to comprehensive assessment of all known human mutations that manifest progressive neurological disease^6^ and establishment of the first precisely (CRISPR Cas9) edited, humanized NKH mouse model expressing a prevalent disease mutation ^19^ with potential to measure post-natal brain disease in mice.

Gene therapy presents potential for curative therapies across a wide clinical spectrum of genetic disorders: the most recent tools are AAV vectors engineered from a non-pathogenic, naturally replication-deficient parvovirus ^20, 21^. Crossing the blood brain barrier (BBB) presents a significant challenge in targeting systemically administered gene therapy to the brain. However, specific serotypes like AAV9 cross the BBB allowing access to the central nervous system (CNS) after systemic administration ^22^. Notably, AAV9 gene therapy received FDA approval for the treatment of Spinal Muscular Atrophy 1 (SMA 1), proving effectiveness of an AAV9 recombinant in crossing the BBB in treatment of human disease ^23^. Here, we provide the first evidence that novel AAV9 recombinants of GFP and GLDC administered systemically in mice, can also access the brain. Along with benefitting the liver, we show that rAAV9-GLDC effects unexpected mechanisms that promisingly treat post-natal astrocyte loss and neurogenesis as well as long term neurological disease and sudden death observed in an advanced preclinical NKH mouse model.

## Materials and Methods

### Animals

A mouse model of attenuated NKH was developed by using CRISPR to engineer the A394V point mutation in *Gldc* in the C57BL/6 mouse strain by Jackson Labs (Bar Harbor, ME, USA) [20]. Homozygous mutant *Gldc*^(A394V/A394V)^ mice were obtained by heterozygous intercross breeding. Mice were housed in Freimann Life Sciences Center at the University of Notre Dame under controlled conditions, with 55% humidity, temperatures of 20-22°C, a 12-hour light-dark cycle with light from 7am to 7pm, and food and water *ad libitum*. Mutant *Gldc^A^*^394^*^V/A^*^394^*^V^*mice treated with the rAAV9 vector were housed singly or in same-sex pairs in a BSL-2 facility. Untreated mutant *Gldc^A^*^394^*^V/A^*^394^*^V^*mice, heterozygous *Gldc^A^*^394^*^V/+^*mice, and wildtype *Gldc^+/+^*mice were housed in cages of 2-5 mice of the same sex. All mouse models and experiments were approved by the Institutional Animal Care and Use Committee of the University of Notre Dame (IACUC 24-01-8316).

### Genotyping

Genotyping was undertaken as previously described^19^. Briefly, litters of mice were genotyped using 0.5 mm of tissue samples taken from the tail or ear of the mouse between postnatal day 7 and 21 (P7-P21). Tissue digestion was done by boiling tissue at 98°C in 200 μL of 50 mM NaOH for 30 minutes, followed by neutralizing with 20 μL of 1M Tris-Cl (pH 6.8). Phusion Blood Direct PCR kit (Thermo Fisher, Cat#F547L) was used to amplify a 753 base pair segment of *Gldc* that included the A394V mutation site. Specific primers with the following sequences were used, forward: 5’-GTTGCATTTCCGTTTCTGGCT-3’ and reverse: 5’-ACTGCCCTCTTACTTGACCATT-3’. 1 μL of neutralized tissue sample was added to a reaction mix comprising 6.5μL PCR-grade water, 10μL 2X Phusion blood buffer, 1μL of each primer, and 0.5μL of Phusion DNA polymerase. The PCR cycle was as follows: an initial denaturation step at 98°C for 5 minutes; 36 cycles of denaturation at 98°C for 10 seconds, annealing at 62°C for 15 seconds, and elongation at 72°C for 30 seconds; and a final elongation at 72°C for 10 minutes. Following amplification, the PCR product using 0.5 μL of Hpy166II restriction enzyme and 2μL of CutSmart buffer (NEB, Cat# R0616S) was incubated at 37°C for one hour. The resulting DNA fragment bands were separated via agarose gel electrophoresis on a 2.5% agarose gel, run at 125V for 30 minutes. Bands were visualized by ultraviolet radiation using a GelDoc 1000 gel imager (BioRad, Part# 400-0065). Wild-type DNA resolves as a single band at 753 bp, whereas DNA containing the homozygous mutation splits the DNA into two bands at 569 and 184 bp. Therefore, a distinct genotype could be determined for each mouse.

### Vector construction, production, and injection

The rAAV9 vector designed for use in this study was prepared at the University of Pennsylvania, Perelman School of Medicine Gene Therapy Vector Core. The vector encoded a 5.1 kb segment of *Mus musculus Gldc* DNA with a CAG promoter, a hybrid construct that fuses the cytomegalovirus enhancer with the chicken beta-actin promoter. The vector was generated by triple transfection of HEK293 cells with a *cis*-plasmid containing *Gldc* and AAV9 ITRs, a *trans*-plasmid encoding AAV9 *rep* and *cap* genes, and an adenovirus helper plasmid. The vector was purified by tangential flow filtration and iodixanol gradient. Vector titer was measured by qPCR. The vector was suspended in sterile phosphate-buffered saline with 5% glycerol and stored in 100μL aliquots of 1.82×10^13^ vector genomes/mL at -80°C. In an early experiment, ten mutant *Gldc^A^*^394^*^V/A^*^394^*^V^* mice between the ages of P25-P36 were injected with 1.0×10^12^ vector genomes of rAAV9-*Gldc* for analyses of liver and blood. In subsequent studies dedicated to brain analyses, a total of 86 mutant mice were each injected intraperitoneally with 1.0×10^12^ vector genomes of rAAV9-GLDC (42 mice) or rAAV9-GFP (44 mice), balanced for gender. Five WT mice were injected with rAAV9-GFP. Intraperitoneal delivery was utilized due to the large number of animals needed for these studies.

### Plasma isolation

Approximately 20 μL of blood was collected from mice via cheek bleed monthly. The blood was collected in heparinized tubes and stored on ice to prevent coagulation. The blood was centrifuged at 1500 x g for 10 minutes at 4°C and the plasma supernatant was isolated. The supernatant was centrifuged at 14000 RPM for 10 minutes at 4°C and the plasma supernatant was isolated and stored at -80°C.

### Glycine quantification

Plasma glycine levels were quantified using a fluorometric glycine assay kit from Sigma-Aldrich (Cat# MAK26), Abcam (Cat# ab211100), or BioVision (Cat# K589-100). All three kits had identical reagents and instructions. We followed instructions without modification provided by the manufacturers. In short, we used the standard curve generated, RFU values were converted to glycine concentrations in nmol/well. The dilution factor (50X) was considered, and glycine concentrations were converted from nmol/well to nmol/μL. Mean glycine values and standard deviations were calculated for each sample and used in the analysis.

### Western Blots

Hemisected mouse brain tissue was lysed with tissue protein extraction reagent (TPER) (*Thermo Fisher Scientific*, 78501) and complete protease inhibitors (1 tablet/10 mL, *Roche*, 11836170001). Homogenized brain extracts were processed further via sonification using 5 seconds of pulsing and 5 seconds of rest for 2 rounds at 40 mA. The samples were then spun down at 14,000 rpms at 4°C in 25 minutes. Protein concentration was quantified with a bicinchoninic acid assay (BCA, *Thermo Fisher Scientific,* 23225). Lysates were separated by SDS-PAGE at 150V for 65 minutes on a 10% acrylamide gel and transferred to a PVDF membrane at 100V for 75 minutes. Membranes were blocked with 5% non-fat milk in 1X Tris-buffered saline with tween (TBST) for 1 hour at room temperature. Primary antibodies in 1x 5% non-fat milk were added to the membrane and probed overnight at 4°C. A secondary antibody in 1x 5% non-fat milk was added and kept on a rocker for 1 hour at room temperature. Membranes were then washed (3x) off once again in 1X TBST and imaged with Pierce enhanced chemiluminescence reagents (700 µL, Thermo Fisher Scientific, 32106) and exposed using Hyperfilm ECL film (*Cytivia*, 28906839). Primary antibodies used were as follows: anti-GLDC (rabbit, Thermo Fisher, PA5-22101, 1:800), anti-GFAP (mouse, Thermo Fisher, 14-9892-82, 1:2000), anti-Iba1 (rabbit, Wako, 016-20001, 1:500), anti-Vinculin (mouse, Millipore, V9131, 1:4000), anti-GFP (mouse, Santa Cruz Biotechnology, SC-9996, 1:600), anti-NeuN (mouse, Thermo Fisher, MA5-33101, 1:500), anti-CNPase (rabbit, Thermo Fisher, PA5-27972, 1:2000), anti-β3-Tubulin (mouse, Thermo Fisher, MA1-19187, 1:2000), anti-C3 Complement (rabbit, Thermo Fisher, PA5-21349, 1:1000), and anti-Lipocalin2 (mouse, Biotechne, AF1857, 1:500). Secondary antibodies were goat anti-mouse IgG (Bio-Rad, 1706516, 1:5000) and goat anti-rabbit IgG (Bio-Rad, 1706515, 1:5000).

### Long term neurological disease, death (LND) and ventriculomegaly (VM)

Starting at 2 months of age, all mouse groups were exposed to transient pain stimuli (of cheek bleed pricking) and subsequently assessed for the adverse events of rearing, falling over and barrel rolling along the head-tail axis, lasting for 1.5-3 min, as measured by the 5-point Racine scale ^24^ and death. Sudden death (described as demise for no apparent reason after they had been alive 24 hours earlier) was also recorded. VM detected in brain sections of mice at one and five months was quantified using ImageJ ^25^ by examining the lateral ventricular volume in coronal sections of brains immune-stained for the glial fibrillary acidic protein (GFAP). The cerebral lateral ventricular volume was quantified by measuring the ventricular area in each hemisphere. The sum of the area across sections was multiplied with slice thickness (35 μm) to obtain the total volume.

### Cytokine analysis

Brain samples from wild type and NKH mice treated with rAAV9-GFP or rAAV9-GLDC, were flash frozen at -80°C until the time of assay. Before cytokine analysis, organs were homogenized in an MP FastPrep tissue homogenizer for 5 minutes. Homogenates centrifuged at 11,000 x *g* for 10 minutes Brain supernatants were probed for IL-1α, IL-1β, IL-2, IL-3, IL-4, IL-5, IL-6, IL-9, IL-10, IL-12 (p40), IL-12 (p70), IL-13, IL-17A, Eotaxin, G-CSF, GM-CSF, IFN-γ, KC, MCP-1 (MCAF), MIP1-α, MIP-1β, RANTES, TNF-α, TGF-β1, TGF-β2, and TGF-β3 using Bio-Plex Assay kits (cat#: M60009RDPD and 171W5001M) from Bio-Rad Laboratories following the manufacturer’s protocols.

### Statistical analysis

Statistical analysis was performed in GraphPad Prism (version 9.4.1) for both glycine analyses and western blots. The mean glycine level ± SD was calculated for glycine comparisons among treatment groups, and a two-way ANOVA with genotype and time as factors, followed by Tukey’s multiple comparisons test, was performed. For within-subject glycine comparisons, a two-tailed, paired Student’s t-test was used. A comparison of protein quantification among treatment groups was performed using one-way ANOVA. A p-value of <0.05 was considered significant (*). In studies of LND and VM, the Mann-Whitney test was used to calculate p values (* < 0.05).

## Results

### Characterization of humanized pA394V NKH mouse model and initial assessment of the effects of rAAV9-GLDC in plasma- and liver-responses

Our large-scale genotype-phenotype studies of 255 GLDC mutations predicted that the prevalent human p.A389V mutation would yield the highest level of attenuated disease ^4, 6^. CRISPR Cas9 was used to edit the equivalent murine mutation p.A394V with expectation to capture at least subset of disease symptoms and lifespan effects in laboratory mice^19^. Since gene therapy is expected to deliver a single effective dose in the life of a patient, we planned long-term studies beginning with investigating baseline characteristics of the p.A394V mouse colony for over 1 year.

As summarized in **Fig. 1A**, in a colony of 1000 mice, homozygous mutants, heterozygotes and wild type mice showed birth rates of 16.7%, 52% and 31% respectively. Approximately 9% of prenatal deaths and 32% of post-natal deaths were due to hydrocephaly (**Fig. 1 A-B**). Relative to wild type, the mutants showed a small reduction in weight at 3 weeks, but that difference reduced and was eliminated by 4-5 weeks (**Fig. 1D**). Virtually, no deaths were detected after 20 weeks (or 4.5 months; **Fig 1C**). This suggested that therapies could be evaluated for their preventable effects on NKH disease phenotypes and death, seen up to 5 months. For the gene therapy studies described here, we injected mice at one month (between the ages of 4-5 weeks), the youngest group found to be substantially free of hydrocephaly.

**Figure 1.**
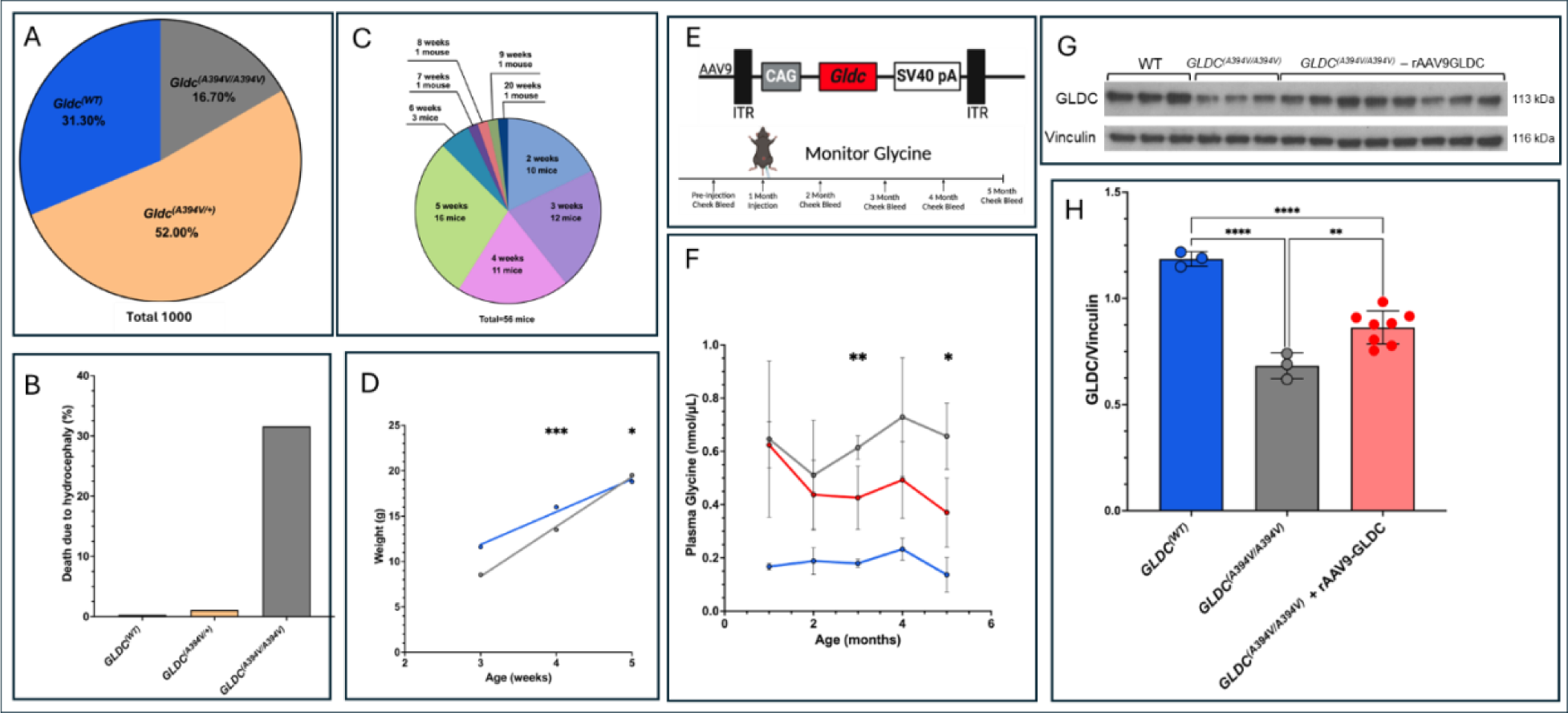
Characterization of the *GLDC* p.A394V mutation mouse colony and effects of administration of rAAV9-GLDC detected in plasma and liver. **A**. Yield of mouse progeny (%) from mating heterozygous *GLDC* ^(A394V/+)^ over one year. Offspring distribution was 31% wild type (blue), 52% heterozygotes (orange, *GLDC* ^(A394V/+)^) and 16.7% homozygous mutants (grey, *GLDC* ^(A394V/A394V)^). **B**. Percentage mortality due to hydrocephaly for each mouse genotype shown in A, namely wildtype (blue), heterozygous (orange) and mutant (grey). **C**. Death due to hydrocephaly was seen 56 of 177 mutant *GLDC* ^(A394V/A394V)^ progeny mice. **D.** Simple linear regression of average weights of *GLDC* ^(A394V/A394V)^ and *GLDC* ^(A394V/+)^ genotypes at 3, 4, and 5 weeks of age. **E**. Schematic of viral vector and the timeline of gene therapy administration and response assessments. **F.** Plasma glycine levels in wild type (blue line), mutant *GLDC* ^(A394V/A394V)^ untreated (black line) and mutant *GLDC*^(A394V/A394V)^ + rAAV9-GLDC treated (red line) mice. **G**. Gender balanced examination of GLDC protein levels in wild type, heterozygous *GLDC* ^(A394V/+)^ and mutant *GLDC* ^(A394V/A394V)^ + rAAV9-GLDC. The band at 113 kDa indicates GLDC protein and the band at 117 kDa indicates the vinculin loading control. **H**. Quantification of the western blot shown in panel G; wild type (n=3 females), untreated mutant *GLDC* ^(A394V/A394V)^ (n= 3 females) mice, and mutant *GLDC^(A^*^394^*^V/A^*^394^*^V)^* + rAAV9-GLDC (n=8, 5 females and 3 males). Data shown in **G-H** are from duplicate experiments.

We selected the AAV9 viral vector because it is known to cross the BBB and enter the CNS ^26, 27^ as well as sustain expression in non-CNS tissues. A novel AAV9 recombinant with a copy of wild-type murine *Gldc* DNA driven by an enhanced constitutive promoter was delivered by intraperitoneal injection to mice **(Fig 1E)**. To assess long term biological efficacy *in vivo*, treated mice were examined for effects on systemic metabolic disease by tracking plasma glycine levels, elevation of which is a high predictor of NKH ^10^. As shown in **Fig. 1F**, in wild type mice, plasma glycine levels remained consistently low (0.2 nmoles/μl) over five months. In mutants *Gldc*^(A394V/A394V)^ prior to treatment, plasma glycine levels were elevated and ranged from 0.4 to 0.8 nmoles/μl. rAAV9-GLDC treatment had no measurable effect on plasma glycine at two months. However, by three months, rAA9-GLDC significantly reduced glycine levels, and in five month old mice, the average plasma glycine levels reduced to 0.4 nmoles/μl. Together, these data suggested that a single injection of rAAV9-GLDC at one month was sufficient to significantly reduce glycine at five months and implicated that functional rAAV9-GLDC was delivered in a sustained manner to *Gldc*^(A394V/A394V)^ mutants to reduce systemic metabolic disease.

Long term expression of GLDC protein in the liver was also evaluated since the organ is known to have an active GCS ^28^ that may impact metabolite levels in plasma. Notably, livers of mutants exhibited a significant reduction in levels of GLDC when compared with wild type mice (**Fig. 1 G-H**). Treatment with rAAV9-GLDC conferred a 25-50% increase in GLDC protein compared to untreated counterparts. The increased levels of GLDC in the liver at nine to ten months are expected to influence long term reduction of plasma glycine levels in circulation.

### rAAV9 accesses the brain in the *GLDC*^(A394V/A394V)^ NKH mouse model

Since AAV9 does not integrate into the genome, the data from **Fig. 1** validated the biological activity of the vector rAAV9-GLDC as well as its capacity to persist in a stable, extrachromosomal, functional state for a substantial fraction of the mouse lifespan. This suggested rAAV9-GLDC may also be useful to query the consequences of long-term effects of gene therapy in other organs, such as the brain, known to be profoundly affected by this neurometabolic disease, but whose cellular disease pathologies and their responsiveness to gene therapy remain completely unknown. However, to treat neurological features and brain pathologies of clinical NKH, systemically administered gene therapy must access the brain.

Since the brain is inarguably complex, we began understanding the effects of the p.A394V mutation on the whole organ. We first evaluated GLDC levels in the brain of wild type and mutant mice at one month of age to establish the extent of loss of protein, prior to the administration of gene therapy. As shown in **Fig 2A-B**, both male and female mutant mouse brains showed ∼85% reduction in GLDC protein, suggesting they were highly depleted compared to wild type counterparts. To these mice, we administered rAAV9-GLDC or a ‘mock’ treatment of rAAV9-Green Fluorescent Protein (GFP) and evaluated long-term effects up to five months (**Fig. 2C**). We used rAAV9-GFP to assess varying efficiencies of the AVV9 vector to deliver cargo to the brain and as a vehicle control for non-specific effects due to GFP protein expression unrelated to GCS activity of GLDC. In anticipation of heterogenous responses arising from indicated vector effects as well as disease manifestation intrinsic to NKH, we placed forty-two mice in the rAAV9-GLDC group and another forty-four in the rAAV9-GFP group balanced for gender (as described in Materials and Methods). In both groups, we delivered equivalent ranges of 0.5 to 1.5 x 10^8^ vector genome copy (vgc)/mg body weight **(Fig. 2D**; also see Methods and Materials).

**Figure 2.**
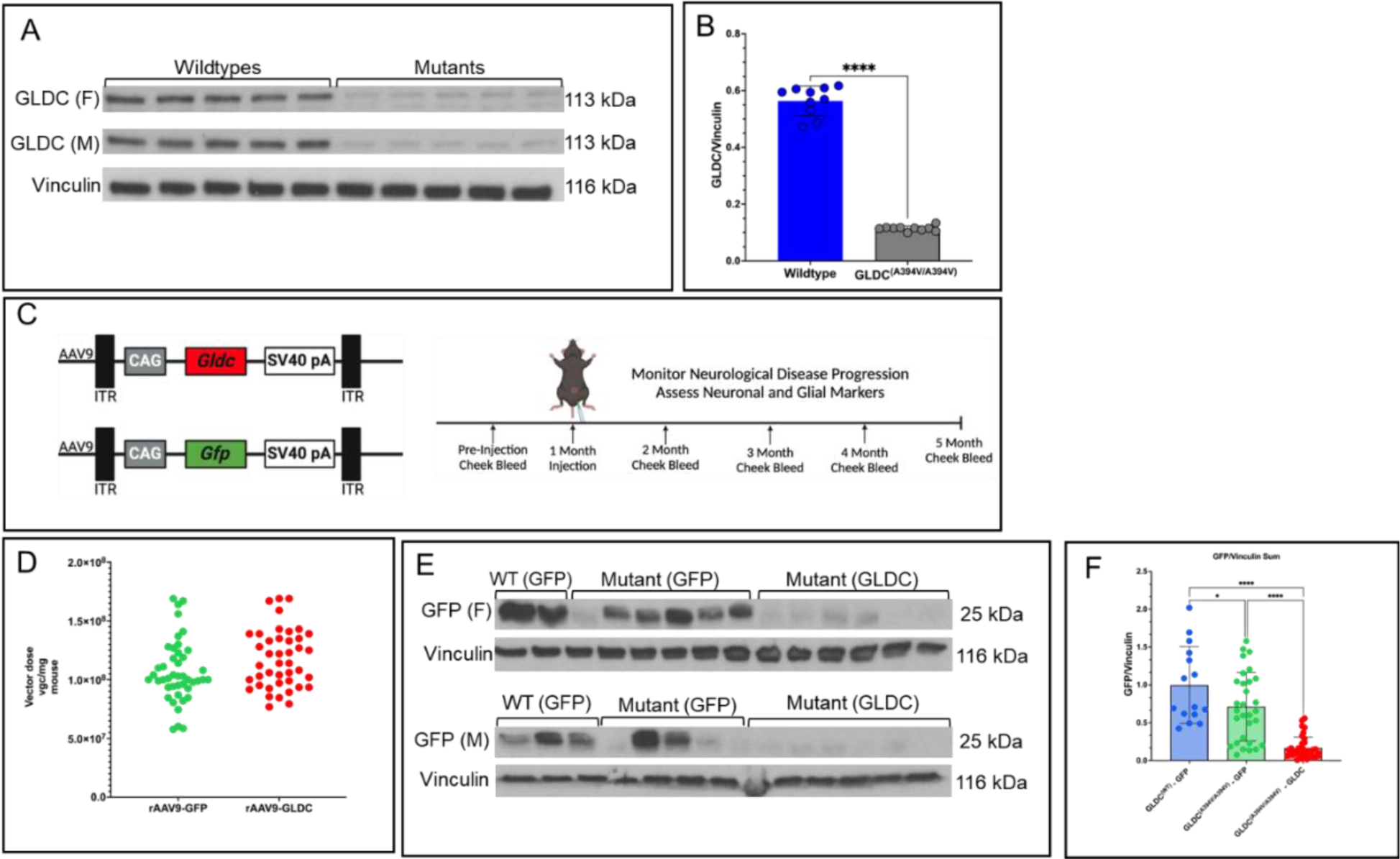
Administration of AAV9 recombinants to *GLDC*^(A394V/A394V)^ mice at one month of age, monitored up to five months. **A-B.** Whole brain, western blots and associated quantification of GLDC protein detected in one-month old wild type and *GLDC* ^(A394V/A394V)^ mutant male (M) and female (F) mice, prior to administration of viral vectors. Vinculin was the loading control. **C.** Representation of the rAAV9 vector containing either *Gldc* or *Gfp* gene and experimental timeline of pre- and post-injection monitoring of mice. **D.** Single dose (vgc/mg mouse) plot comparing intraperitoneal rAAV9-GFP (green) or rAAV9-GLDC (red) respectively delivered to 44 and 42 *GLDC* ^(A394V/A394V)^ mice (balanced for gender). **E-F.** Whole brain western blots and quantitation of GFP detected in a gender balanced group of five-month-old wild type and *GLDC* ^(A394V/A394V)^ mice that received a single injection of rAAV9-GFP at one month.

Western blots were undertaken to assess whether systemic administration with ‘mock’ treatment using rAAV9-GFP construct at one month of age resulted in detectable GFP levels in the brain at five months. As shown in **Fig. 2 E-F**, we found that GFP was detected in the brains of a majority of mutants as well as wild type mice. Specifically, in eleven randomly selected mutants, GFP was found in brains of five of six females and three of five males, suggesting robust brain access and persistence in >70% of injected mice (**Fig. 2 E-F**). Further, the lack of detectable GFP in the remaining ∼28% may reflect less efficient delivery beyond the sensitivity of western blots (rather than failure to deliver any protein to the brain). Since GFP was also detected in the brain of wild type mice (**Fig. 2 E-F**), our novel recombinant AAV9-mediated delivery of cargo across the blood-brain barrier was not due to a special characteristic of mutant mice.

### Effects of gene therapy on glial cells in the NKH brain

Clinical studies suggest that although the corpus callosum is prominently affected in NKH, other major regions of the brain are also affected ^29^. The human protein atlas also indicates high expression of GLDC throughout the brain^13^ suggesting need for analyses of whole brain. We began with evaluating astrocytes since GLDC is prominently associated with these major glial populations in the brain ^14, 15^, particularly in the mouse brain (brainrnaseq.org). We assessed levels of astrocytes at five months of age in the whole brain in mice that received either rAAV9-GFP or rAAV9-GLDC (at one month of age) by measuring GFAP a pan-astrocytic biomarker, via western blots analysis. As shown in **Fig. 3A-B**, GFAP levels were significantly lower (by 17%) in the *GLDC*^(A394V/A394V)^-GFP group compared to their wild type counterparts. Yet, notably, at one month old mutants showed no change in GFAP levels compared to wild type counterparts (**Fig. 3C**). In rAAV9-GLDC mice at five months, we saw a significant (35%) increase of GFAP relative to age-matched rAAV9-GFP (with measurable increase in both genders **Fig. 3A-Bi-iii**). This reflected an 18% GFAP increase even when compared to WT. Overall, these data suggest that between 1 and 5 months of age astrocyte levels declined in mutant animals. However, introducing a wildtype copy of GLDC promoted astrocyte production in mutants that exceeded levels seen in wild type brains (**Fig. 3B**).

**Figure 3.**
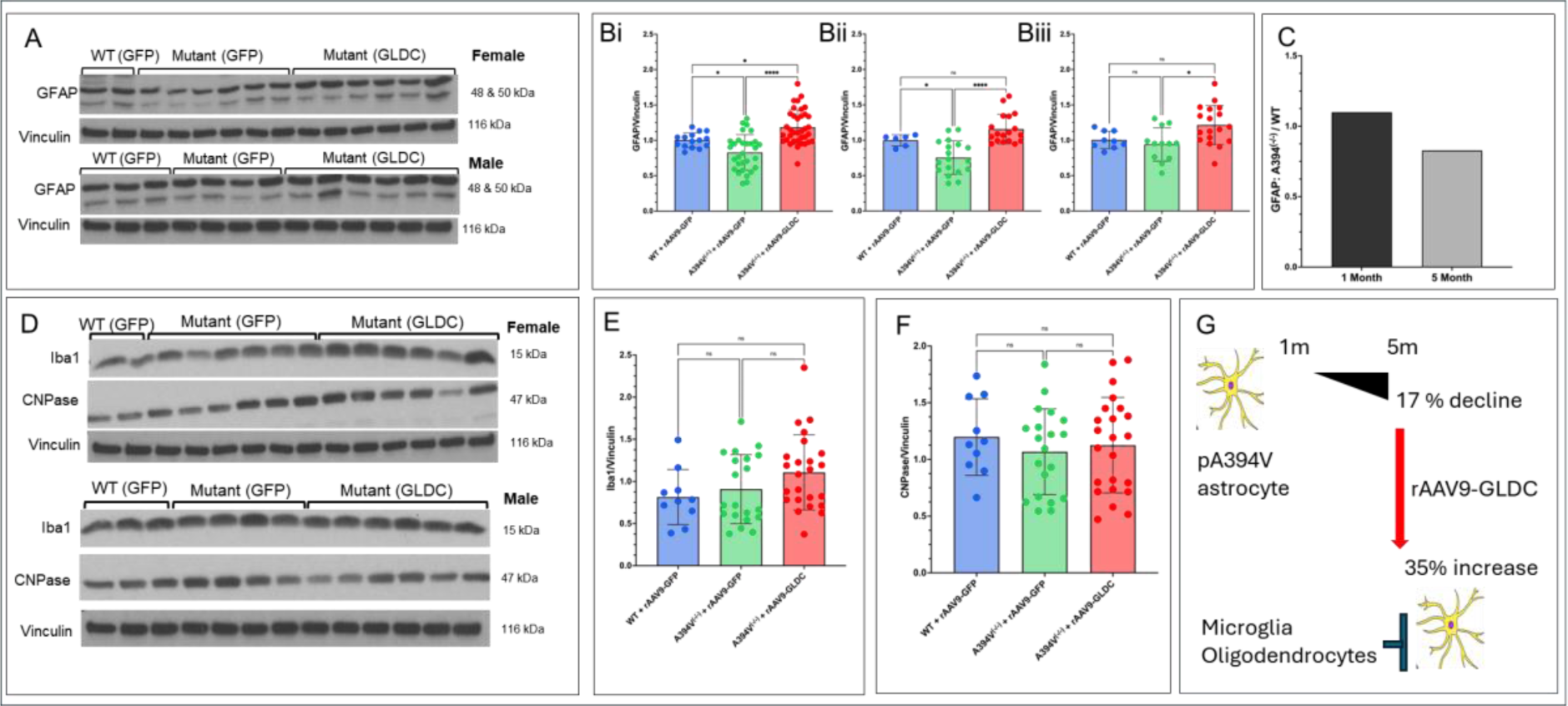
Effect of rAAV9-GLDC and rAAV9-GFP on glial cell proliferation in NKH mutants. **A-B.** Representative, whole brain, western blots and associated quantitation of astrocyte levels using GFAP as a pan-astrocytic biomarker in male and female wild type and mutant mice. Vinculin was used as the loading control. **Bi** combines data from females (**Bii**) and males (**Bii**i) of wild type (blue bar), rAAV9-GFP treated-*GLDC* ^(A394V/A394V)^ (green bar), and rAAV9-GLDC treated-*GLDC*^(A394V/A394V)^ (red bar) mice. **C.** Ratio of GFAP-positive astrocytes in mutants relative to wild type mice at one month (black bar) and five months (grey bar) of age. **D-F.** Representative whole brain western blots and associated quantitation of glial markers Iba1 (for microglia) and CNPase (for oligodendrocytes). **E-F** shows combined male and female data for wild type (blue bar), rAAV9-GFP treated-*GLDC* ^(A394V/A394V)^(green bar) and rAAV9-GLDC treated-*GLDC* ^(A394V/A394V)^ (red bar) mice. **G.** Proposed model of astrocyte proliferation following rAAV9-GLDC treatment in mutant *GLDC*^(A394V/A394V)^ mice without activation of microglia or oligodendrocytes.

To assess additional glial cells, we examined microglia, which in neurodegenerative diseases may typically be the first to activate and initiate an inflammatory cascade that triggers astrocyte activation and proliferation ^30^. However, as shown in **Fig. 3D-E**, compared with wild type, microglia levels (detected by tracking ionized calcium binding adaptor molecule 1; Iba1) showed no significant changes in mutants treated with either rAAV9-GFP or rAAV9-GLDC. Since the corpus callosum is a major white matter region commonly affected in NKH clinical disease, we further evaluated oligodendrocyte expression to assess white matter degeneration. Oligodendrocyte activation was measured by quantifying the levels of 2’,3’-cyclic nucleotide 3’-phosphodiesterase (CNPase), a major enzyme marker for oligodendrocyte activation ^31^. Brain CNPase levels did not show statistically significant differences between wild type, mutants treated with rAAV9-GFP, or mutants treated with rAAV9-GLDC (**Fig. 3D and 3F).** The collective findings on glial cells are summarized in a model shown in **Fig. 3G**, which highlights that early quantitative reduction in GLDC is associated with reduction of astrocytes (but not microglia or oligodendrocytes) as detected in the brains of five-month-old NKH mice. Administration of rAAV9-GLDC boosts astrocytes without activation of microglia or oligodendrocytes, suggesting alternative mechanisms of astrocyte proliferation are induced by gene therapy in the NKH brain.

### Effects of gene therapy on astrocyte inflammation and neurogenesis

In response to injury, astrocytes proliferate by a process called astrogliosis ^32^, which activates inflammatory pathways that can be measured by activation of complement C3, a primary signal that begins the inflammatory cascade ^33, 34^. As shown in **Fig. 4A-B** mutant brains presented a small reduction of complement C3, but this change was not statistically significant when compared to responses of wild type mice. Administration of rAAV9-GLDC to mutants, similarly, only slightly raised C3, to levels seen in wild type mice. As an additional marker, we used lipocalin 2 (LCN2) as a second measure of astrocyte activation. It has been shown that proinflammatory astrocytes express high levels of LCN2 reflecting inflammation driven by the transcription factor nuclear factor kappa B (NF-kB) ^35^. Conversely, in anti-inflammatory astrocytes, LCN2 expression is minimal ^36^. Our analyses suggested that relative to wild type, LCN2 was significantly decreased in NKH mutant injected with rAA9-GFP and a further decrease was seen in mutants injected with rAAV9-GLDC (**Fig. 4A, C**). Together, the data in **Fig. 4A-C** strongly suggest that astrocyte proliferation stimulated by gene therapy does not induce a proinflammatory response and is likely to occur independent of astrogliosis.

**Figure 4.**
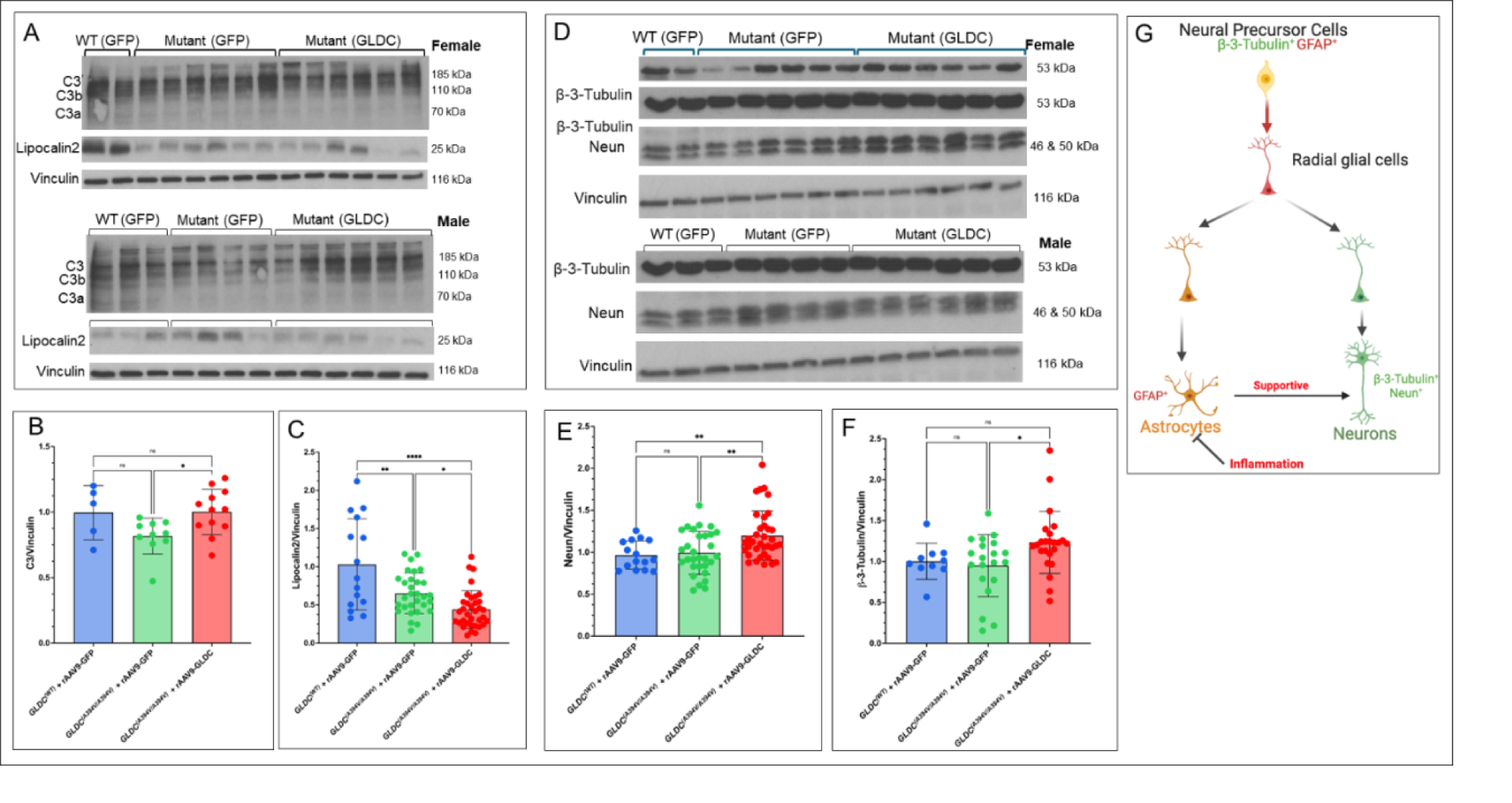
Effect of rAAV9-GLDC and rAAV9-GFP on astrocyte inflammation and neurogenesis. **A-C.** Representative, whole brain western blots and quantitation of (**A-B**) complement C3 and (**A,C**) lipocalin 2 (LCN2) in both male and female mice. **D-F.** Representative, whole brain, western blots and quantitation of (**D-E**) mature neurons (NeuN) and (**D, F**) neural precursors (β-3-tubulin). Vinculin was the loading control. **G.** Model showing rAAV9-GLDC induces stimulation of neural precursors through pathways of neurogenesis and astrogenesis resulting in production of both neurons and astrocytes (which may further support neurons) in absence of inflammation. The model contrasts astrogliosis which is accompanied by pro-inflammatory glial cascades seen in most neurodegenerative diseases.

Low expression of LCN2 has been suggested to foster astrocytes that create a neuroprotective environment ^36^ at the subgranular zone (SGZ) and subventricular zone (SVZ) ^37^. These astrocytes develop by promoting the proliferation and differentiation of neural precursor or neural progenitor cells, undergoing astrogenesis in the absence of inflammation, a process that may occur concomitantly with neurogenesis ^38^ ^39^. We therefore evaluated effects on neurogenesis by examining levels of mature neurons as well as markers of neural precursor cells by measuring the levels of NeuN (hexaribonucleotide binding protein-3), a marker for mature neurons as well as β-3-Tubulin (Tubb3) that marks neural precursors. As shown in **Fig. 4D-F**, we found that administration of rAAV9-GLDC (but not rAAV9-GFP) increased levels of NeuN and β-3-tubulin. This strongly supports that rAAV9-GLDC acts by stimulating the proliferation of neural precursor or neural progenitor cells to promote astrogenesis and neurogenesis as proposed in the model in **Fig. 4G**. Consistently, GLDC has been shown to regulate the maintenance and induction of pluripotency as well determine the fate of stem cells ^40, 41^. We additionally propose that newly formed and proliferating astrocytes may provide additional support to neurons, further improving brain function after gene therapy (**Fig. 4G**).

### Long term neurological disease, sudden death and cerebral atrophy are prevented by administration of rAAV9-GLDC

Clinical NKH presents complex neurological disease and brain pathologies that result in death. In mouse models, adverse neurological responses can be enhanced by external stimuli^42^ We observed consequences subsequent to transient pain stimulus of cheek pricking that all mice were subjected to in monthly bleeds, after two months of age. These included convulsive events lasting for 1.5-3 min (measured by the 5-point Racine scale) ^24^ as well as sudden death (as described in Materials and Methods). As shown in **Fig 5A**, 29.5% (13 of 44) GFP-mice presented adverse events that we cumulatively characterize as long-term neurological disease and death (LND) seen up to five months of age. Four of the thirteen responded to cheek pricking with immediate convulsions and/or death. Eight underwent sudden death and one mouse died of developing hydrocephaly at nineteen weeks (4.5 months). During this period, astrocytes were the only major cell type in the brain reduced in mutant brains (while levels of microglia, oligodendrocytes and neurons were unchanged compared to wild type: see Fig. 3 and 4). LND events were entirely prevented in mutants administered with rAAV9-GLDC (**Fig. 5B)**, which had astrocytes and neurons boosted (as previously shown in Figs 3-4), Together these data suggested astrogenesis and concomitant neurogenesis induced by gene therapy, protected against severe and fatal disease progression.

**Figure 5.**
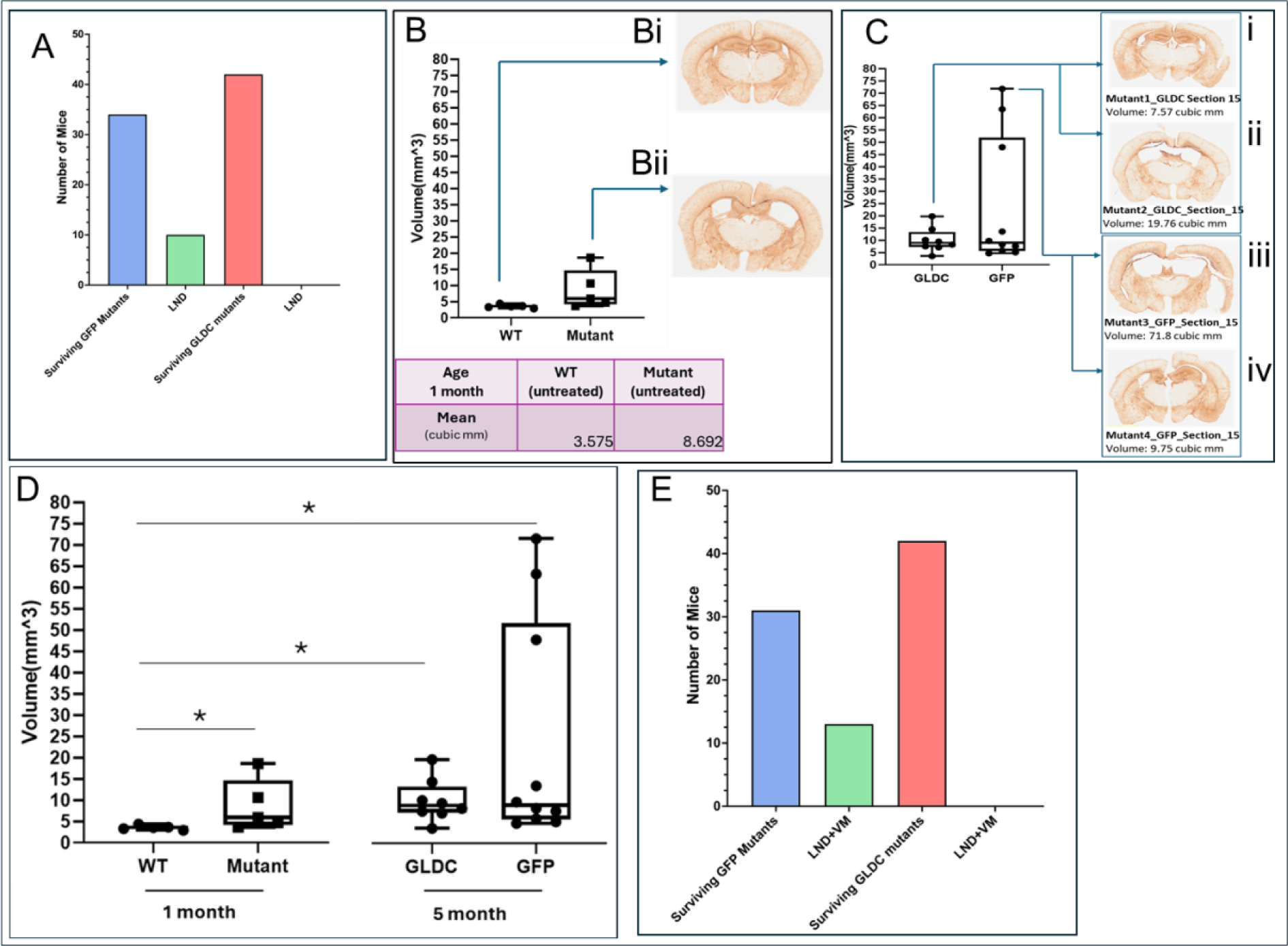
Protective effect of gene therapy on post-natal long term neurological disease and death (LND) and ventriculomegaly (VM). **A**. Mutant *GLDC* ^(A394V/A394V)^ mice were ‘mock’ treated with a single dose of rAAV9-GFP. Of 44 mice, 29.5% showed symptoms of LND (stimulus-induced convulsions and death) over five months (green bar). Of 42 mutants administered a single dose of rAAV9-GLDC (red bar), none develop LND. **B.** Protective effect of gene therapy on post-natal VM in mutant mice. Five mutant (non-hydrocephalic) mice *GLDC* ^(A394V/A394V)^ at one month of age showed VM with a mean 2.5-fold volume increase of ventricles relative to wild type (WT) counterparts. **Bi-ii** Pathologies of representative sections. **C.** VM was evaluated in *GLDC* ^(A394V/A394V)^ mutants after administration of rAAV9-GFP (10 mice) or rAAV9-GLDC (7 mice). **Ci-iv** show examples of VM pathologies seen in coronal brain histopathology of equivalent sections. **D.** Direct comparison of ventricular volumes measured in panels B and C. **E.** Mutant *GLDC* ^(A394V/A394V)^ ‘mock’ treated with a single dose of rAAV9-GFP showed LND+VM in 37% of mice (green bar) Mutant *GLDC* ^(A394V/A394V)^ that received a single dose of rAAV9-GLDC (red bar) failed to develop LND or VM progression.

We also examined VM, a disease condition where brain ventricles become enlarged. It is distinguished from hydrocephalus, where the enlargement occurs due to increased pressure from cerebrospinal fluid (CSF)^43^ . Both events are seen in patients with NKH ^5, 8^. In p.A394V mutant mice, hydrocephalus developed in the first three weeks, but VM is caused by brain dysgenesis or atrophy that occurred over a longer period of several months and could not be distinguished by external doming. In **Figs. 5 B-C**, we provide examples of coronal brain histopathology and range of ventriculomegaly pathologies seen in equivalent sections of wild type and mutant *GLDC*^(A394V/A394V)^ mice at one month of age as well as mutants treated with rAAV9-GLDC or rAAV9-GFP at five months. As summarized in **Fig. 5D**, *GLDC*^(A394V/A394V)^ mutants (that were non-hydrocephalic) at one month of age showed a mean 2.5-fold volume increase of ventricles relative to their wild type counterparts. A subset of mutant *GLDC*^(A394V/A394V)^ mice that were mock treated by administration with a single dose of rAAV9-GFP and examined at five months of age showed a considerable worsening of VM. In contrast, *GLDC*^(A394V/A394V)^ that were administered rAAV9-GLDC did not worsen. A total of seventeen mutant mice in five months were randomly selected and used in pursuing outcomes with rAAV9-GFP (10 mice) and rAAV9-GLDC (7 mice) and compared to five wild type and five mutants at one month of age. When the total LND counts were added to confirmed cases of VM, 37% of mock-treated mice were shown to progress with the disease and death, while mice administered with rAAV9-GLDC were completely protected (**Fig 5E)**. These data combined with our findings from **Fig. 4**, suggested that rAAV9-GLDC gene therapy has the potential for treating neurological disease, cerebral atrophy and death arising from loss of astrocytes and neurons.

### Implication of NKH and rAAV9 therapy for inflammatory processes in the brain

While glycine accumulation in the brain is known to disrupt neurotransmitter balance and lead to neurological disorders such as seizures, it’s unclear whether this disease process also triggers brain inflammation. Furthermore, it is not known whether loss of astrocytes in mutants from one to five months is associated with inflammation. In this context, the inflammatory response to rAAV9-GFP and rAAV9-GLDC therapies is unknown. To investigate, we measured and compared the levels of 26 cytokines in three groups: wild-type mice treated with rAAV9-GFP, *GLDC(^A^*^394^*^V/A^*^394^*^V^)* mice treated with rAAV9-GFP, and *GLDC(^A^*^394^*^V/A^*^394^*^V^)* mice treated with rAAV9-GLDC. Notably, the majority of cytokine levels were significantly lower in the *GLDC(^A^*^394^*^V/A^*^394^*^V^)* mutant mice compared to wild-type mice treated with rAAV9-GFP (**Fig. 6A-D**), with the exceptions of IL-1β, IL-4, IL-17, GM-CSF, TGF-β2, and TGF-β3 (**Fig. 6C**). Notably, IL-1α, IL-2, Il-3, I-5, IL-13, Eotaxin, G-CSF, KC, MCP-1, MIP-1β, and TNF-α were significantly decreased in rAAV9-GLDC-treated *GLDC(^A^*^394^*^V/A^*^394^*^V^)* mutant mice when compared to rAAV9-GFP-treated wild-type mice (**Fig. 6D**). Interestingly, MIP-1α levels were restored to near wild-type levels in AAV9-GLDC-treated *GLDC(^A^*^394^*^V/A^*^394^*^V^)* mutant mice (**Fig. 6E**). While the primary functions of MIP-1α are associated with inflammation and immune responses, some studies indicate that it can play a complex and multifaceted role in neurogenesis^44, 45^ ^46^. This result correlates with our findings in **Fig. 4** and **5**, where AAV9-GLDC treatment promotes neurogenesis.

**Figure 6.**
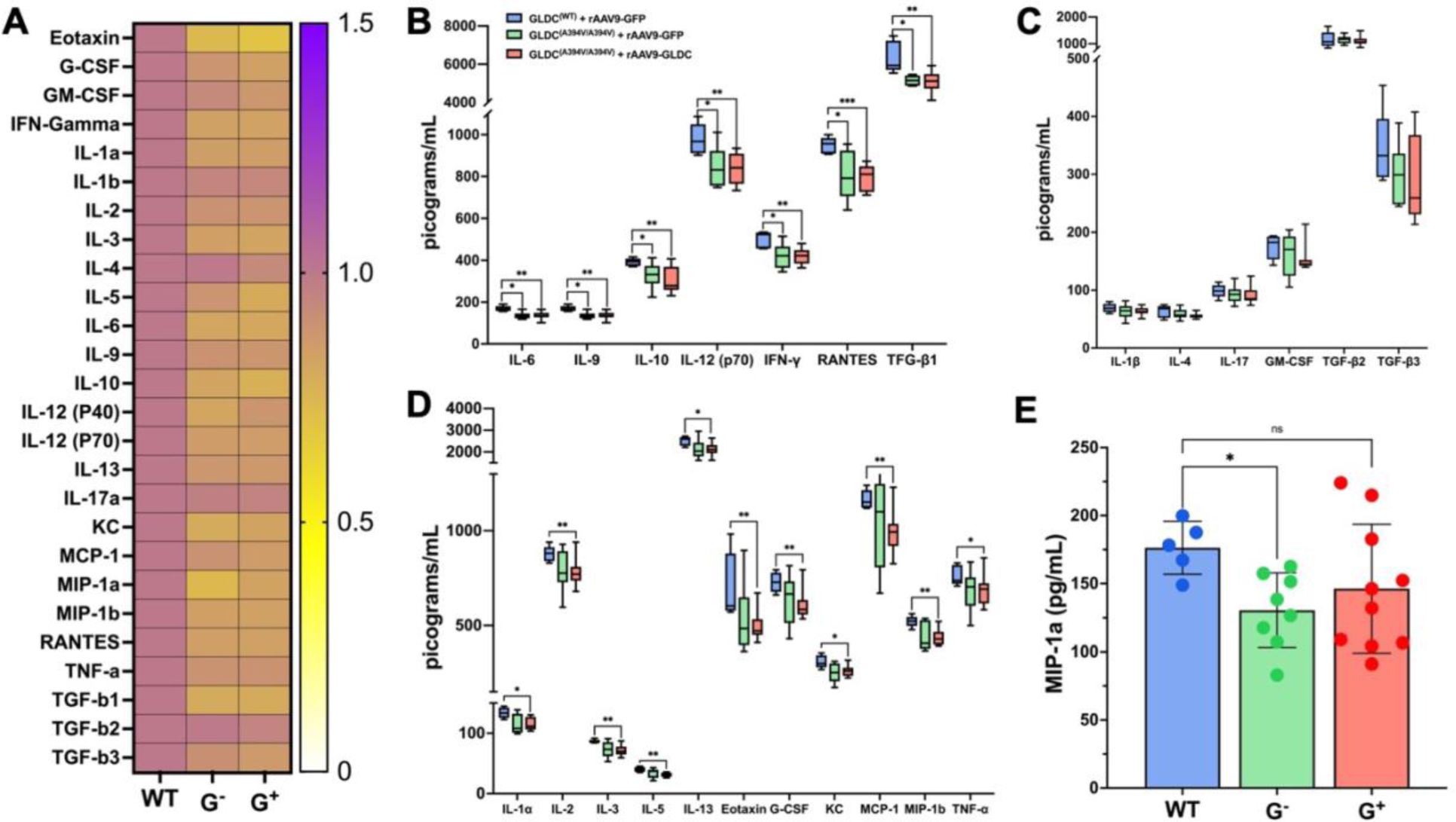
Brain inflammatory responses in mice with defect in GLDC and/or administration of rAAV9 vectors. **A**. Heatmap represents cytokine fold changes in the brain of GLDC^-/-^ mutant mice administered with rAAV9-GFP (G^-^; green) or rAAV9-GLDC (G^+^; red) over wild type treated with rAAV9-GFP (WT; blue) brains. **B**. Cytokines that were significantly decreased in GLDC^-/-^ mutant mice regardless of the treatment. **C.** Cytokines that were significantly decreased only in GLDC^-/-^ mutant mice treated with rAAV9-GLDC. **D**. MIP-1α brain levels in GLDC-/- mutant mice treated with rAAV9-GLDC are rescued to wild type mice. The horizontal bar represents the median value with range as error bars. The Mann-Whitney U test was used; *, P < 0.05 was considered statistically significant. **, P < 0.005.

## Discussion

Our studies provide the first demonstration that systemic administration of a novel rAAV9 recombinant can access systemic organs like the liver as well as the brain to confer therapeutic benefits and protect against disease and death in NKH mice. Prior studies suggest that an AAV9 vector failed to access the brain in a gene trap (null) NKH mouse model ^47^. Our work differs since we utilized a stronger expression construct, a humanized mouse mutation model, and assessed long-term benefits over many months, because all three features are highly relevant to future patient treatment. Thus, our novel AAV9 recombinant, where gene expression is driven by a chicken β-actin promoter and the enhancer elements of cytomegalovirus (CMV), elicited beneficial responses in the liver and enabled delivery of gene therapy across the BBB in a prevalent patient mutation mouse model. In this study we focused largely on the brain, since its cellular and pathological disease correlates and their corrections are expected to be critical to understanding and treatment of NKH. There was heterogeneity in the delivery of protein cargo to the brain. However, over 70% of mice showed a substantial presence of GFP in the brain after systemic delivery of 0.5 to 1.5 x 10^14^ vgc/kg, which is within the range of low to moderate doses used in human gene therapy studies ^23^. In patients, we expect intravenous administration in the head region would be an appropriate delivery mechanism which has been successfully used for brain delivery of therapeutic AAV9 vectors in treating spinal muscular atrophy ^23^.

Our studies show that all NKH mutant mice treated post-natal with rAAV9-GLDC at one month of age, were reduced in metabolic and cerebral disease and death as measured up to five months. Mice mature much faster than humans and their maturation does not correlate linearly with human aging. Current assessment suggests that mice, at one month, are close to fourteen years of human age, and at six and ten months close to thirty-four and fifty years of human age, respectively. Although these age conversions remain estimates, they suggest that our novel rAAV9-GLDC recombinant may be effective in treatments initiated even in teenage years and approach decades of protective benefit. As with all gene replacement therapy, administration at young ages (particularly for severe mutations) is expected to confer the highest level of preventative benefit.

The principal effect of the prevalent mutation in our humanized mouse model was the destabilization of GLDC, which over five months mainly led to the reduction of astrocytes but not oligodendrocytes, microglia, or neurons. Astrocytes are known to express high levels of GLDC, and in our model, over the first five months may play an important role underlying the major disease symptoms. However, mutant brains show lower levels of inflammatory cytokines, suggesting defective astrocytes and their loss, are not associated with induction of inflammatory processes in the brain. Our novel vector rAAV9-GLDC induces proliferation of astrocytes in absence of inflammatory cues of microglia or oligodendrocytes. Inflammatory process intrinsic to astrocytes were also not elevated. In particular, that TNFα, IL1β and IL6 do not increase, strongly support that astrocyte proliferation is not due to inflammatory activation of microglia^30^. Rather astrocyte proliferation was associated with neurogenesis suggesting involvement of neural precursors. Recent studies with human induced pluripotent stem cells (iPSCs) from an NKH patient suggest deficiency in *Gldc* in neural progenitor cells influences astrocyte and neuronal lineage ^17^. This suggests that GLDC plays a key role in differentiation and proliferation of neural progenitor cells which is consistent with our model that rAAV9-GLDC complements a neural progenitor defect in GLDC to support renewal and replenishment of astrocytes and neurons in the brain (**Fig. 4G**). In this context it is interesting to note that the corpus callosum (white matter tract) known to be affected in clinical NKH has recently been shown to promote astrogenesis ^48^.

NKH presents as a complex neurological, muscular, and respiratory disease in patients. Over 50 distinct symptoms have been reported, including hypotonia, seizures, gait imbalance, dysphagia, and autism, among others. Although seizures are particularly prominent in severe cases, no single clinical symptom is shared across all patients^6^. Capturing disease symptoms such as seizures has posed a challenge in NKH mouse models.^49^. Our humanized mouse model exhibited post-natal death due to hydrocephalus within 3-4 weeks after birth, similar to prior studies^49^. In addition, mice that did not receive rAAV9-GLDC remained fragile during the subsequent period of 1 to 5 months, during which 37% of these mice showed adverse effects without overt hydrocephalus: ∼ two-thirds of which experienced ‘sudden death’ and one-third exhibited convulsions or death in response to transient painful stimuli. Due to their fragility, we did not attempt to monitor electrical seizure activity, as this is an invasive process that could lead to death and/or severe infections. A few mice showed VM, and one mouse at 4.5 months demonstrated hydrocephalus. When considered collectively, there was a stark difference between the adverse events experienced by GFP-mice compared to none in the GLDC-mice. We could not pinpoint the cause of many adverse events (such as sudden death), but all were protected by rAA9-GLDC. Marker analyses confirmed that adverse events were not due to inflammation in the brain. Inflammation, particularly of the liver, can be elicited by AAV gene therapy^23^. However, the lack of GLDC expression in microglia and macrophages (brainrnaseq.org) suggests that inflammation may not present a major risk for NKH gene therapy.

## Acknowledgments

We thank Vaishnavi Reghunath (IISER Pune) for Image J analyses. We thank the members of the Haldar lab for helpful discussion during the course of the work.

## Animal facility

We thank the Freimann Life Sciences Center (FLSC) and staff at the University of Notre Dame, for providing excellent services for mouse housing, caring, breeding, blood drawing, overall monitoring and veterinarian support.

## Penn Vector Core

We thank the Penn Vector Core for producing rAAV9-GLDC and rAAV9-GLDC.

## Regulatory Approval

All procedures were approved by the Institutional Biosafety Committee (IBC), University of Notre Dame. Design of animal studies and procedures regarding their use were approved by the Institutional Animal Care and Use Committee (IACUC) University of Notre Dame.

## Author Contribution

Alejandro Lopez-Ramirez – conceptual design, experimentation, analyses and interpretation of studies in Fig. 1, 2, 3, 4, 5, 6 and 7, injections, colony management organ harvest, visualization of results, writing of related methods and figure legends, abstract, introduction, results, discussion and overall primary drafting and editing of the manuscript.

**Figure 7.**
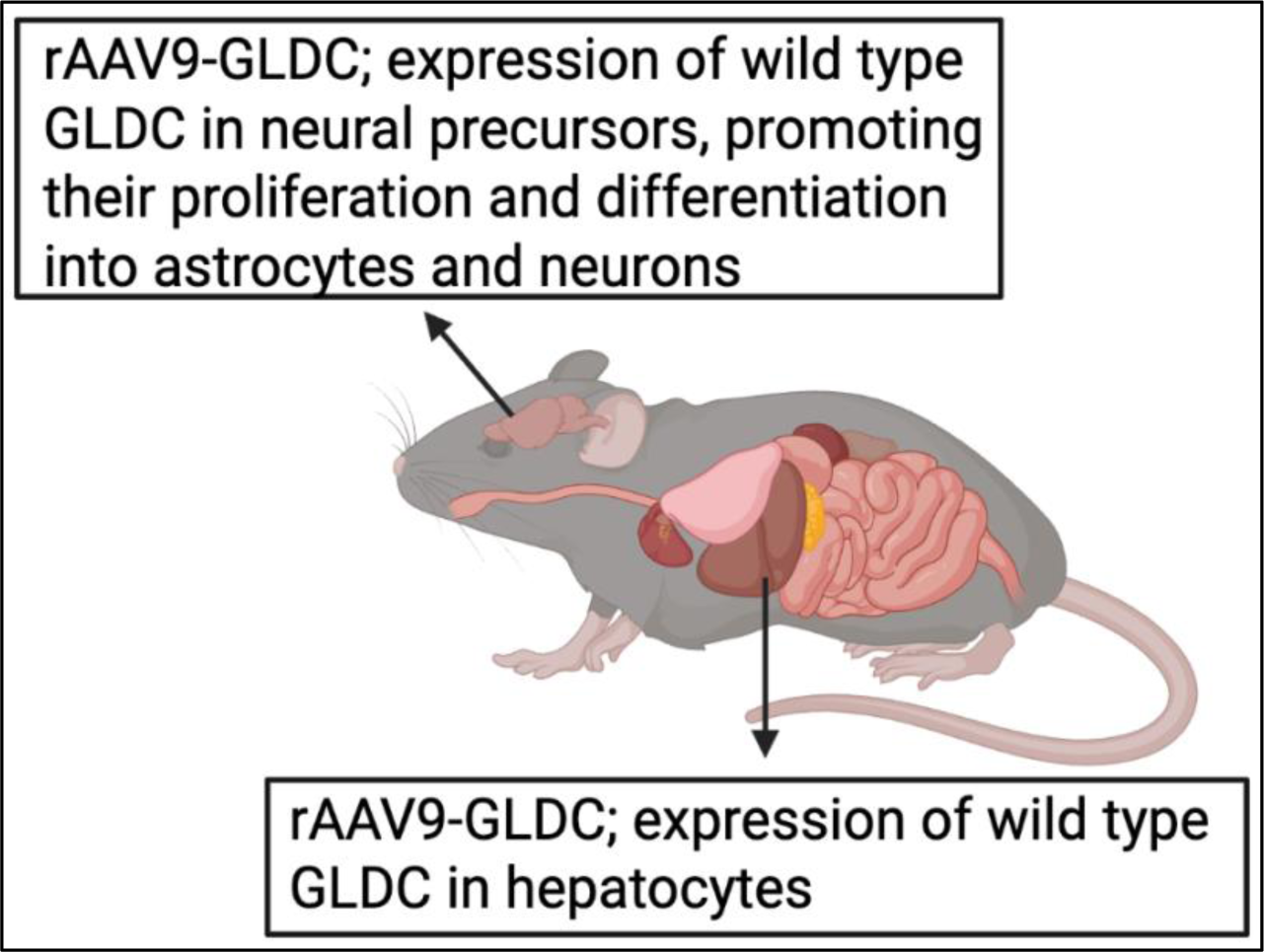
Summary of mechanisms of novel AAV9-GLDC gene therapy to treat the brain and liver in a humanized point mutation mouse model of NKH. The model summarizes the effects of single intraperitoneal administration of rAAV9-GLDC that can target both the brain and liver of *GLDC*^(A934V/394V)^ mutants. In the brain, proliferation of astrocytes linked to neurogenesis (through stimulation of neural precursors) confers protection against long term neurological disease and brain pathology, in absence of inflammation. In the liver, the observed elevation of GLDC in hepatocytes is expected to significantly reduce levels of blood glycine. In summary, a single dose effects normalization of NKH disease and death, is protective over many months, suggesting long term benefits expected of gene therapy is expected to be a vast improvement over current treatment for NKH.

Adviti Bali – conceptual design, experimentation, analyses and interpretation of studies in Fig. 1, 2, 5 and 6 (and relevant supplementary data), colony management organ harvest, visualization of results, writing of related methods and figure legends, abstract, introduction, results, discussion and overall primary drafting and editing of the manuscript.

Md Suhail Alam – vector design and construction, conceptual design, experimentation, analyses of Figure 1 and relevant supplementary data, injections, colony management organ harvest and editing the manuscript.

Prasad Padmanabhan – conceptualization, design experimentation and analyses of Figure 2 and relevant supplementary data, colony management, injections, organ harvest, and editing of the manuscript.

Shaun Calhoun – experimentation and analyses of Figure 5 and relevant supplementary data, colony management, injections, organ harvest and editing of manuscript.

Caroline Bickerton - experimentation, analyses and interpretation of studies in Fig. 1 and editing the manuscript.

Ana L. Flores-Mireles design and interpretation of inflammation data (Fig 3 and Fig. 6) and editing the manuscript.

Kasturi Haldar – overall conceptualization, design and supervision of entire project, design and development of vectors, all data analyses and visualization of results, primary drafting and editing of manuscript, funding acquisition.

